# Phytochemical profiling, antioxidant capacity, and in vivo safety assessment of Warrigal spinach (*Tetragonia tetragonioides*) and Kensington Pride mango (*Mangifera indica*) extracts using Zebrafish larvae

**DOI:** 10.64898/2026.04.27.720960

**Authors:** Sarah Kiloni, Akhtar Ali, Frank Dunshea, Jeremy Cottrell, Paolin Rocio Cáceres-Vélez, Patricia Regina Jusuf

## Abstract

Plant-derived bioactive compounds are recognised for their antioxidant potential and benefit for human diseases, including age-related diseases caused by oxidative stress. However, their antioxidant composition and safety profiles remain insufficiently understood. This study integrates phytochemical profiling, antioxidant evaluation, and *in vivo* toxicological assessment of Warrigal spinach (*Tetragonia tetragonioides*) and Kensington Pride mango (*Mangifera indica*). Spinach exhibited greater antioxidant capacity, and higher total phenolic and flavonoid content than mango: TPC (14.2 ± 0.6 mg GAE/g vs 1.30 ± 0.07 mg GAE/g) and TFC (9.61 ± 0.39 mg QE/g vs 0.08 ± 0.0 mg QE/g). LC-ESI-QTOF-MS/MS identified 187 metabolites dominated by flavonoids (53.5%) and phenolic acids (16%), with spinach showing greater chemical diversity. Quantitative analysis revealed higher levels of hydroxycinnamic acids and flavonoid glycosides in spinach, whereas mango contained distinct metabolites, including mangiferin and pyrogallol. Zebrafish embryo / larval assays demonstrated high safety margins, with LC_50_ values of 478.8 mg/L (spinach) and >480 mg/L (mango). At 480 mg/L spinach displayed developmental abnormalities and malformations. These findings demonstrate that antioxidant capacity is linked to phenolic composition, but does not predict toxicity. Thus, integrated phytochemical and safety evaluation for extracts with complex compound mixtures are critical to identify botanicals suitable for future drug development.

**Highlights:**

- Warrigal spinach exhibited >10-fold higher phenolic content and antioxidant capacity than Kensington Pride mango.
- LC-ESI-QTOF-MS/MS identified 187 metabolites, with flavonoids as the dominant phytochemical class.
- Zebrafish assays confirmed high safety margins, demonstrating no direct correlation between antioxidant capacity and toxicity.

Graphical abstract

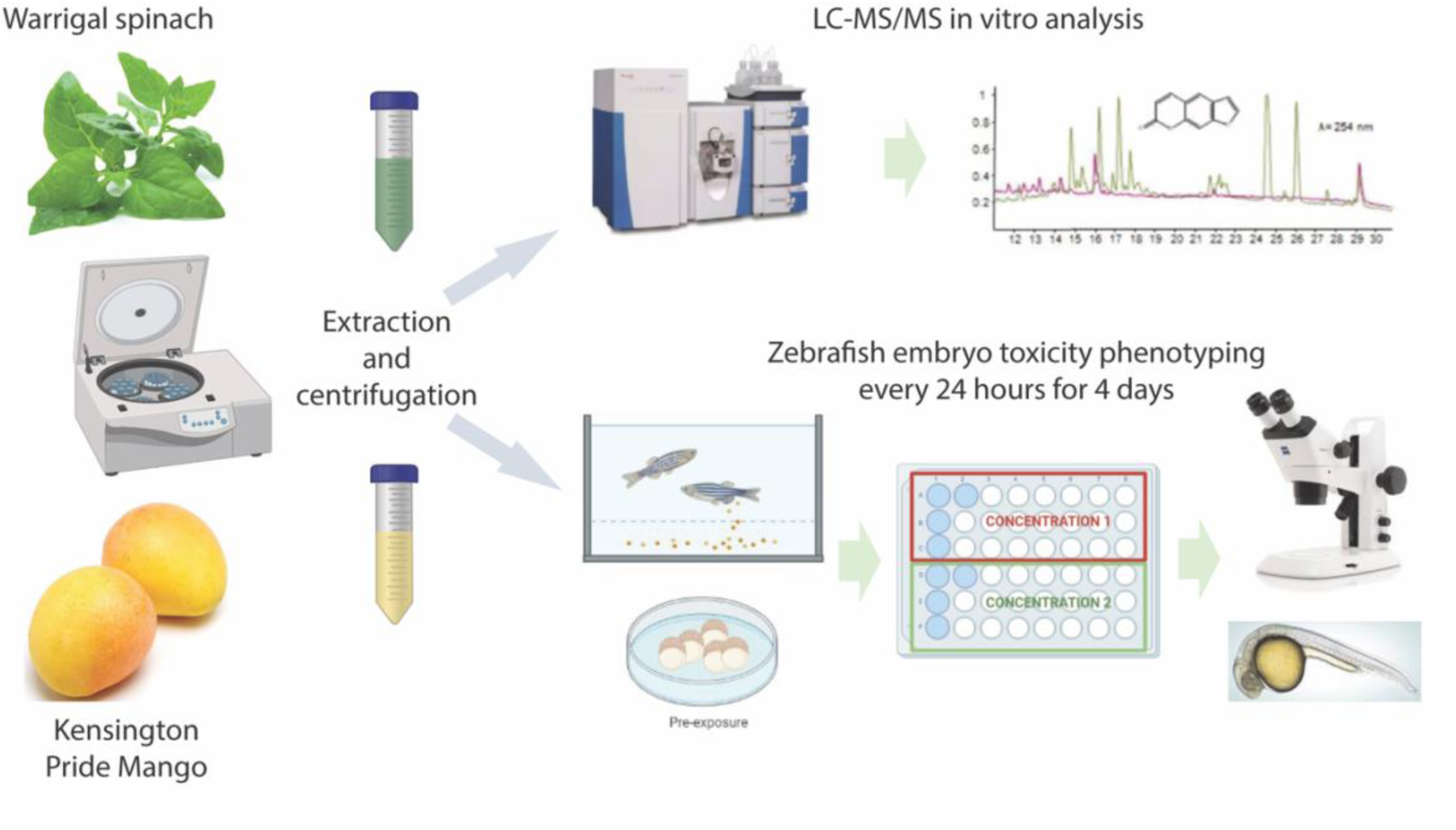

## 1. Introduction

Oxidative stress is a hallmark of numerous diseases, including cardiovascular diseases, diabetes, neurodegenerative disorders, and cancer (reviewed in Chandimali et al., 2025; Korovesis et al., 2023; Leyane et al., 2022; Pizzino et al., 2017). Oxidative stress occurs when there is elevated production of Reactive Oxygen Species (ROS), such as occurs in ageing (reviewed in Caceres-Velez et al., 2022; reviewed in Slimen et al., 2014). Excessive ROS accumulation can cause failure to maintain cellular homeostasis and promotes mitochondrial dysfunction, leading to cell damage, oxidation of proteins, lipids and DNA, ultimately promoting cell death (Cross & Templeton, 2004; Gomes et al., 2020; Osborne et al., 2014; Sohal et al., 2000; Zhao et al., 2019; Zorov et al., 2000). Consequently, there has been increasing interest in identifying effective exogenous antioxidants, such as phytochemicals, flavonoids, carotenoids, vitamins, polyphenols, and phenolic acids, particularly those derived from plant-based sources (reviewed in Chandimali et al., 2025; Korovesis et al., 2023; Pizzino et al., 2017).

Plant-derived bioactive compounds, especially polyphenols and flavonoids (Jayaprakasha et al., 2000; Simioni et al., 2018), are widely recognised for their potent antioxidant properties (Tugen & Buruleanu, 2025) and remarkable potential for targeting pathologies of human disorders. Antioxidant rich native plants from different geographical regions (e.g. Southeast Asia) have been cultivated for centuries and are being actively investigated for toxicological *in vivo* effects (Dissanayake et al., 2022; He et al., 2018; Ismail et al., 2017; Ong et al., 2015; Zhang et al., 2023). Australia is home to a diverse range of native and cultivated plant species, which have been an important part of Indigenous history, culture and medicinal applications, and have been explored for their antioxidant, anti-inflammatory and antimicrobial properties (Ali, Cottrell, et al., 2022; Akhtar Ali et al., 2023; Ali et al., 2024; Cartwright, 1991; Rupesinghe et al., 2016; Tan et al., 2011a, 2011b). Despite the growing body of research on their bioactive composition, there is a limited understanding of the relationships among phytochemical profiles, antioxidant capacity, and the safety of these plant-derived extracts (Mani et al., 2021). This represents a major gap that requires a high throughput standardized model pipeline that allows direct correlation of bioactive compound combinations with safety assessment, and comparison between studies.

Warrigal spinach (*Tetragonia tetragonioides*), a native leafy vegetable of Australia and New Zealand, has historically been used as a dietary source of nutrients and antioxidants (Lee et al., 2018). It is known to contain high levels of vitamin C, carotenoids, and phenolic compounds, which contribute to its potential health benefits (Dissanayake et al., 2022). On the other hand, Kensington Pride mango (*Mangifera indica*), one of the most widely cultivated mango varieties in Australia, is recognised for its rich content of bioactive compounds such as mangiferin, carotenoids, and phenolic acids (Joyce et al., 2002; Rocha Ribeiro et al., 2007). Although both plants possess antioxidant properties, their phytochemical composition and functional potential differ significantly due to variations in plant tissue type, metabolism, and environmental exposure.

Previous studies have largely focused on either phytochemical characterisation or antioxidant activity of plant extracts, often without integrating toxicological evaluation. However, the assumption that natural compounds are inherently safe can be misleading, as phytochemicals may exhibit cytotoxic, genotoxic, or developmental effects depending on their concentration and interactions within complex mixtures (Ali, Kiloni, et al., 2022). Therefore, evaluating both the efficacy and safety of plant-derived compounds is essential for their potential application in the food and pharmaceutical industries. The zebrafish (*Danio rerio*) embryonic model has emerged as a powerful *in vivo* system for high-throughput toxicological screening. Zebrafish are small, freshwater aquatic organism which are ideal for genetic, toxicological, developmental, and biomedical research (Chahardehi et al., 2020; Falcao et al., 2018; Garcia et al., 2016; Hermsen et al., 2011; Lim et al., 2022; Lima et al., 2025), due to their genetic similarity to humans, rapid development, and transparency (Howe et al., 2013). The use of zebrafish embryos and larvae as a model organism for toxicity assays (FET – fish embryo acuate toxicity test) to assess mortality and morphological parameters, has increased significantly during the last decade (Ali, Kiloni, et al., 2022; Cáceres-Vélez et al., 2022; Ghorbani et al., 2022; Lima et al., 2025; Tanguay, 2025). Zebrafish provide an important bridge between *in vitro* assays and mammalian models, and allow for simultaneous assessment of biological activity and safety (Cáceres-Vélez et al., 2022).

In this study, we aimed to evaluate the antioxidant capacity, phytochemical composition, and safety profile of Warrigal spinach and Kensington Pride mango extracts. Using a combination of spectrophotometric assays (TPC, TFC, DPPH, ABTS, and FRAP), LC-ESI-QTOF-MS/MS analysis, and zebrafish embryonic toxicity screening, we sought to (i) characterize the bioactive compounds present in these plant extracts, (ii) compare their antioxidant potential, and (iii) assess their safety for potential applications in nutraceutical and pharmaceutical development. This integrative approach provides valuable insights into the relationship between phytochemical composition, antioxidant activity, and biological safety, contributing to the growing field of plant-based functional ingredients.

## 2. Materials and Methods

### 2.1 Plant Extract Preparation

Warrigal spinach leaves and Kensington pride mango fruit were collected and stored at –80℃, thawed prior to extraction and blended to reduce particle size. Extraction of the plants was performed by adding 10 ml of 70% analytical-grade ethanol (Thermo Fisher Scientific Inc., Scoresby, VIC, Australia). Samples were covered with aluminium foil and placed in a shaking incubator at 120 rpm at 4 ℃ for 24 hours. A paper filter was used to remove particles from suspension in the extracts. The solvent from the extracts was removed by lyophilisation. The purified samples were stored at −80℃ and protected from light until use to prevent photo-oxidation of the antioxidants.

### 2.2. Phenolic and Flavonoid Content, and Antioxidant Activity

*In vitro* characterisation was performed as described previously (Ali, Kiloni, et al., 2022; Cáceres-Vélez et al., 2022). The plant extracts were obtained and characterised (polyphenol content, HPLC, and LC-MS) using analytical reagents from Thermo Fisher Scientific Inc. (Scoresby, VIC, Australia), Sigma-Aldrich (Castle Hill, NSW, Australia), and Agilent Technologies (Melbourne, VIC, Australia). Some reagents, such as sodium carbonate anhydrous and hydrogen peroxide (30%), were purchased from Chem-Supply Pty Ltd. (Adelaide, SA, Australia), and sulfuric acid was purchased from RCI Labscan (Rongmuang, Thailand).

#### 2.2.1 Determination of Total Phenolic Content (TPC)

For TPC, 25 μL of each phenolic extract was mixed with 25 μL Folin-Ciocalteu (F-C) reagent and 200 μL Milli-Q water, then incubated for 5 minutes before adding 25 μL 10% sodium carbonate. After 1 hour incubation at room temperature in the dark, the absorbance at 765 nm was recorded. Gallic acid (0-200 µg/ml) was used to generate a standard curve for TPC quantification, and Trolox (0-600 µg/ml) served as the external standard.

#### 2.2.2 Determination of Total Flavonoid Content (TFC)

For TFC, 80 µL of phenolic extract of 80 µL in 96-well plate, followed by shaking after adding 80 µL of aluminium chloride solution (2 %) and 120 µL of sodium acetate (50 g/L). The reaction mixture was then kept in the dark at 25 °C for 2.5 hours. The absorbance was measured at 440 nm, and all samples were analysed in triplicate. The standard curve (0-50 μg/mL of quercetin was generated (R^2^=0.999), and results were presented as mg QE/g.

#### 2.2.3 Determination of Antioxidant Activity

For DPPH, 275 μL of 0.1 mM DPPH dye and 25 μL of the sample were incubated in the dark for 30 minutes, then read spectrophotometrically at 517 nm. DPPH was measured relative to Trolox (0-100 μg/mL). For ABTS, after 16 hours, the absorbance was calibrated to 0.70 ± 0.02, and a 10 μL sample or standard was mixed with 290 μL of ABTS dye in the dark and incubated for 6 minutes at room temperature. A calibration line was obtained by running Trolox (0-200 μg/mL) in methanol. The FRAP reagent was prepared by mixing 300 mM sodium acetate buffer, 10 mM TPTZ, and 20 mM ferric chloride in a 10:1:1 (v/v/v) ratio. Additionally, a 20 µL sample extract was mixed with 280 µL of FRAP reagent in a 96-well plate. The reaction mixture was placed at 37 °C for 10 min, and the absorbance was measured at 593 nm. All results were quantified by constructing a standard curve against 0–50 µg/mL ascorbic acid in water. The results were expressed as mg AAE (ascorbic acid equivalent) /g.

### 2.3 Screening for bioactive compounds in native Australian plant extracts

Polyphenolic compounds were identified following the protocol described by Cáceres-Vélez et al. (2022) using an Agilent 6520 Accurate-Mass QTOF machine. Acquisition (4 spectra/second) was achieved in auto MS/MS positive and negative modes. MassHunter Workstation Software (version B.06.00) was used for the extraction and identification of phenolic compounds. These experiments were completed in duplicate. The quantification of individual compounds was performed using the method with modifications as described by Cáceres-Vélez et al. (2022). The detection wavelengths were set at 280 nm, 320 nm, and 370 nm. The compounds were quantified in triplicate on a dry weight basis (μg/g).

### 2.4 Zebrafish larval acute toxicity tests

#### 2.4.1 Animal husbandry

Zebrafish were maintained at the *danio rerio* (DrUM) facility at the University of Melbourne, Victoria, Australia. All experiments were conducted in accordance with the Code for the Care and Use of Animals in Research and local animal guidelines (ID 22235). For each plant extract, a stock solution was freshly prepared in autoclaved E3 medium (5 mM NaCl; 0.17 mM KCl; 0.33 mM CaCl_2_; 0.33 mM MgSO_4_) and diluted to obtain increasing concentrations (0; 15; 30; 60; 120; 240; and 480 mg/L) for each extract. The pH of extract stock solutions was tested daily and maintained between 6.5 and 8.0. At 3 hpf, zebrafish embryos were exposed in a static system as described (Ali, Kiloni, et al., 2022; Cáceres-Vélez et al., 2022). Briefly, healthy embryos were placed individually into 46-well plates (one embryo/well) containing 500 μL/well of test solution. Phenotypic and behavioural changes were recorded daily (24, 48, 72, and 96 hours of exposure) for each embryo, using a stereomicroscope (LEICA M80). All experiments were conducted in triplicate from different AB WT zebrafish breeding parents (*n* = 60 embryos/concentration) for each plant extract. After phenotyping on the fourth day, larvae were humanely killed (Ehrlich et al., 2019).

### 2.4 Statistical analysis

The data were analysed by one-way analysis of variance (ANOVA) followed by Dunnett’s multiple comparisons test, and a Probit model was fitted to determine the LC_50_. GraphPad Prism 9.4.1 was used, with the significance level set at 5%.

## 3. Results

### 3.1. Measurement of total phenolic contents and their antioxidant activities of Warrigal Spinach and Kensington Pride mango extracts

Warrigal spinach exhibited significantly higher antioxidant capacity than Kensington Pride mango across all evaluated *in vitro* assays (Table 1). The total phenolic content (TPC) of Warrigal spinach was more than 10-fold higher than that of Kensington Pride mango. Similarly, total flavonoid content (TFC) was substantially higher in Warrigal spinach (9.6 ± 0.4 mg QE/g) than in mango, indicating a markedly richer polyphenolic composition. Consistent with these findings, antioxidant assays demonstrated significantly enhanced radical scavenging and reducing capacities in Warrigal spinach. The DPPH assay revealed that Warrigal spinach exhibited more than 11-fold higher radical scavenging activity than Kensington Pride mango. Likewise, ABTS values were approximately 12-fold higher in Warrigal spinach (25.6 mg AAE/g) compared to mango. Although FRAP values were comparatively lower overall, Warrigal spinach still demonstrated approximately 8-fold higher reducing power than Kensington Pride mango (0.13 ± 0.02 mg AAE/g).

**Table 1.** Total phenolic contents and antioxidant activity in Warrigal spinach and Kensington Pride Mango.

| Variables | TPC<br>(mg GAE/g) | TFC<br>(mg QE/g) | DPPH<br>(mg AAE/g) | FRAP<br>(mg AAE/g) | ABTS<br>(mg AAE/g) |
| --- | --- | --- | --- | --- | --- |
| Warrigal Spinach | 14.24 ± 0.61 | 9.61 ± 0.39 | 36.7 ± 1.81 | 1.00 ± 0.21 | 25.6 ± 0.96 |
| Kensington Pride Mango | 1.30 ± 0.07 | 0.08 ± 0.00 | 3.24 ± 0.10 | 0.13 ± 0.02 | 2.04 ± 0.06 |

The results (mean ± standard deviation, n=3) per gram of sample. TPC: Total phenolic content. TFC: Total flavonoid content; DPPH: 2,2'-diphenyl – 1 - picrylhydrazyl assay; FRAP: ferric reducing antioxidant power; ABTS: 2,2'-Azino-bis(3-ethylbenzothiazoline-6-sulfonic acid) assay, GAE: Gallic acid equivalent, QE: Quercetin equivalent, AAE: ascorbic acid equivalent.

### 3.2. LC-MS/MS characterisation and identification of phytochemicals in Warrigal Spinach and Kensington Pride Mango extracts

LC-ESI-QTOF-MS/MS analysis enabled the tentative identification of 187 phytochemicals from spinach (SP) and mango (KP) samples (Table S1 and Figure 1). The detected metabolites belonged to several major chemical classes, including flavonoids, phenolic acids, stilbenes, lignans, xanthones, other phenolic compounds, and non-phenolic metabolites. Among these, flavonoids (Fig. 1A) represented the predominant group, accounting for 53.5% (100 compounds) of the total identified metabolites, followed by other phenols (17.1%), phenolic acids (16.0%), non-phenolic compounds (5.9%), lignans (4.8%), xanthones (1.6%), and stilbenes (1.1%). Further classification of flavonoids revealed eight subclasses (Fig. 1B), with flavones (34%) being the most abundant, followed by isoflavonoids (21%), flavonols (19%), flavanones (13%), chalcones (5%), flavanols (3%), O-methylated flavonoids (3%), and flavans (2%). This distribution highlights the structural diversity of flavonoids present in the analysed fruit samples.

**Figure 1.**
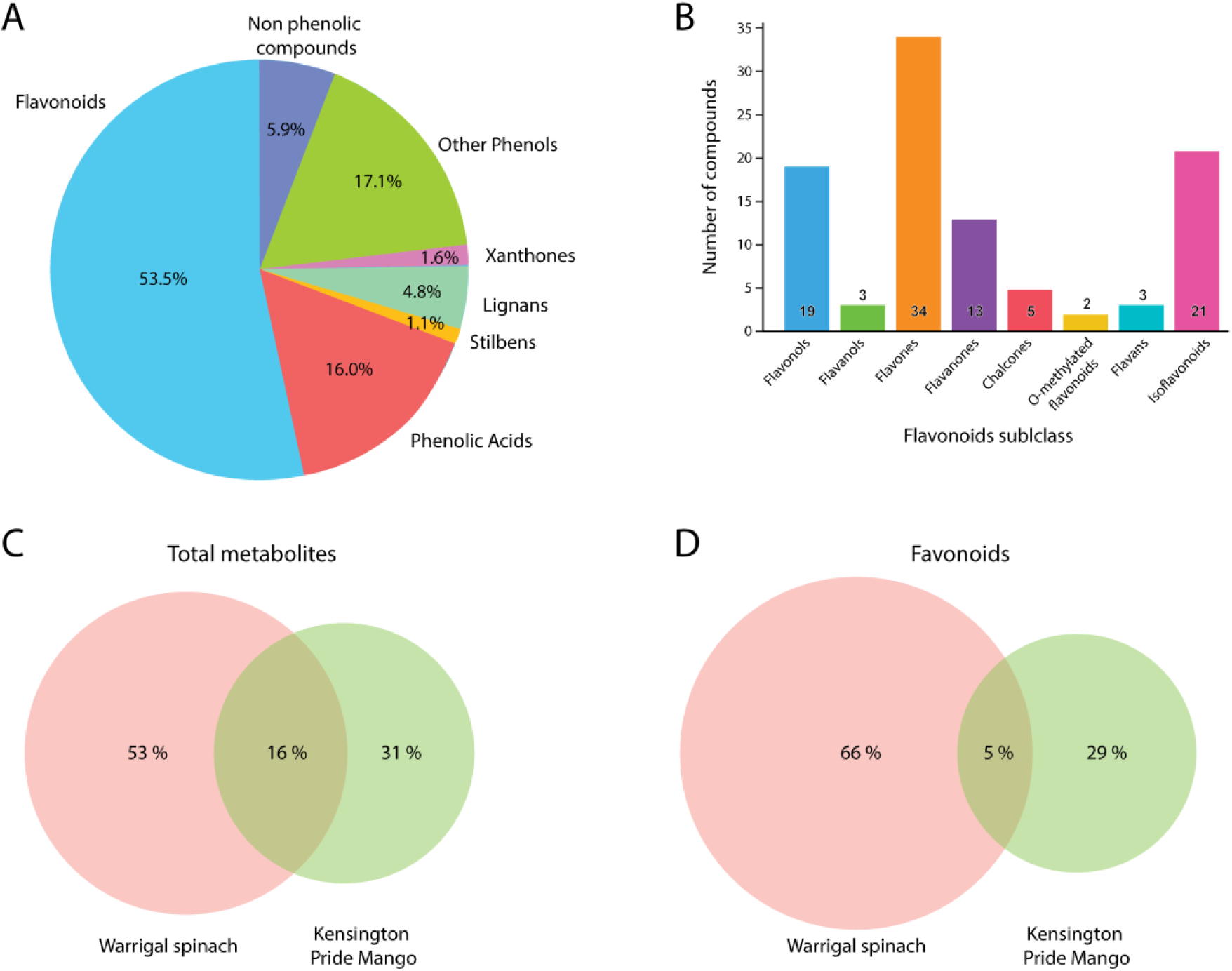
Composition and abundance of phytochemicals identified from LC-ESI-QTOF-MS/MS analysis. (A) Relative composition (%) of main classes of metabolites showed highest abundance of flavonoids. (B) Subclasses of the 100 identified flavonoid compounds, with numbers in bars indicating number of compounds identified. (C, D) Overlap of identified metabolites between Warrigal spinach and Kensington Pride showed quite unique profiles, with only 16% of total metabolites (C) and 5% of all flavonoids overlapping (D).

The Venn diagram analysis of total metabolites indicated a clear difference in phytochemical composition between the two fruits. Approximately 52.9% of the compounds were unique to spinach, 31.0% were unique to mango, while 16.0% were common to both samples (Fig. 1C). A similar comparison of flavonoids showed that 66% of flavonoids were detected exclusively in spinach, 29% exclusively in mango, and 5% were shared between both fruits (Fig. 1D). These results suggest that spinach exhibit greater flavonoid diversity, whereas mango contains a smaller but distinct subset of flavonoid compounds. Overall, the results demonstrate a high chemical diversity in both fruits, with flavonoids being the dominant phytochemical class and spinach contributing the largest proportion of unique metabolites.

#### 1. Phenolic acids

Phenolic acids were primarily detected in negative electrospray ionisation mode as deprotonated molecular ions [M−H]^−^. Their identification was based on accurate mass measurements, retention behaviour, and diagnostic MS/MS fragmentation patterns. Hydroxybenzoic acid derivatives such as gallic acid, protocatechuic acid, and *p*-hydroxybenzoic acid exhibited characteristic product ions associated with decarboxylation (loss of CO_2_, 44 Da). For instance, gallic acid (*m/z* 169) generated a prominent fragment at *m/z* 125, corresponding to the loss of CO_2_ [M−H−44 Da] (Lopez-Herrador et al., 2025). Glycosylated derivatives, including gallic acid 4-*O*-glucoside and protocatechuic acid glucoside, were characterised by the neutral loss of hexose moieties (162 Da), producing aglycone ions typical of hydroxybenzoic acids. Hydroxycinnamic acids such as caffeic acid, ferulic acid, and *p*-coumaric acid displayed typical fragmentation pathways involving decarboxylation and cleavage of side chains. For example, caffeic acid (*m/z* 179) yielded a diagnostic fragment at *m/z* 135 [M−H−44], whereas ferulic acid *m/z* 193) produced characteristic ions at *m/z* 178, 149, and 134 after the loss of CH_3_ and CO_2_ from the precursor and later daughter ions. In addition, quinic acid conjugates such as feruloylquinic acid generated product ions at *m/z* 191 and 173, which correspond to the quinic acid backbone. These fragmentation patterns are widely reported as diagnostic features of hydroxycinnamic acid derivatives (Kiani et al., 2023).

#### 2. Flavonoids

Flavonoids represented the most abundant class of metabolites identified. Structural characterisation of flavonoids relied mainly on glycosidic bond cleavage and retro-Diels–Alder (RDA) fragmentation patterns observed in the MS/MS spectra. Flavonoid glycosides typically exhibited neutral losses corresponding to sugar residues, including 162 Da (hexose), 146 Da (rhamnose), 132 Da (pentose), and 176 Da (glucuronic acid). Flavonol glycosides, such as quercetin and kaempferol derivatives, generated characteristic aglycone ions upon cleavage of glycosidic bonds. For example, quercetin glycosides produced a dominant fragment ion at *m/z* 301, whereas kaempferol derivatives generated an aglycone ion at *m/z* 285. Subsequent fragmentation of these aglycones yielded typical RDA product ions, including *m/z* 179 and 151, which are widely used as diagnostic ions for quercetin-type flavonoids (Kiani et al., 2023).

Flavones and flavanones displayed similar fragmentation behaviour. For example, hesperetin derivatives yielded an aglycone fragment at *m/z* 301, whereas naringenin glycosides produced characteristic ions at *m/z* 271 following glycosidic bond cleavage. Isoflavonoids were identified by heterocyclic ring cleavage and neutral losses of substituents, yielding distinctive fragment ions that supported their structural assignment. Together, these diagnostic fragmentation pathways enabled the tentative identification of multiple flavonoid subclasses, including flavonols, flavones, flavanones, chalcones, flavans, and isoflavonoids (Akhtar Ali et al., 2023; Kiani et al., 2023).

#### 3. Stilbenes and lignans

Stilbenes and lignans were tentatively identified based on their accurate mass values and characteristic fragmentation behavior in negative ion mode. Stilbene derivatives typically exhibited fragmentation involving loss of methyl (15 Da) and methoxy (15 Da) groups, generating product ions associated with the aromatic backbone (Zhang et al., 2019). Lignan compounds, including pinoresinol and sesamin derivatives, exhibited fragmentation patterns arising from ether bond cleavage and demethylation. These compounds frequently produced fragment ions corresponding to losses of CH_3_ (15 Da), CH_2_O (30 Da), and CO_2_ (44 Da). Such fragmentation pathways are typical for lignan structures and provided important evidence for their identification in the analyzed samples (Akhtar Ali et al., 2023).

#### 4. Other compounds

In addition to the major phenolic subclasses, several other phenolic compounds were identified, including arbutin, pyrogallol, quinic acid, and carvacrol derivatives. These compounds exhibited fragmentation pathways involving decarboxylation, demethylation, and cleavage of glycosidic bonds. For example, arbutin ([M−H]^−^ *m/z* 271) generated a fragment corresponding to the loss of a glucose unit (162 Da), yielding the hydroquinone aglycone ion (Ali, Cottrell, et al., 2022). Similarly, quinic acid ([M−H]^−^ *m/z* 191) exhibited characteristic product ions at *m/z* 173 and 127, which are commonly reported as diagnostic fragments for quinic acid derivatives (Ali, Cottrell, et al., 2022). The combined interpretation of accurate mass values and MS/MS fragmentation patterns allowed the tentative assignment of several additional phenolic metabolites in the fruit samples. Besides phenolic metabolites, several non-phenolic secondary metabolites, including terpenoids, steroidal saponins, and related compounds, were also detected. These compounds showed fragmentation patterns characteristic of their structural classes. Terpenoids typically exhibited sequential neutral losses of methyl groups and water molecules, whereas saponins displayed fragmentation resulting from stepwise cleavage of glycosidic linkages. The interpretation of these fragmentation pathways, together with accurate mass data and chromatographic retention behaviour, enabled the tentative identification of the detected non-phenolic metabolites.

### 3.3. Quantification and semi-quantification of phenolic compounds in Warrigal spinach and Kensington Pride mango

The quantification and semi-quantification of selected phenolic compounds in Warrigal spinach and Kensington Pride mango were performed using LC-ESI-QTOF-MS/MS based on peak-area comparison and available reference standards, following previously reported approaches. A total of 16 phenolic compounds were quantified across both samples, including hydroxybenzoic acids, hydroxycinnamic acids, flavonoids, and other phenolic derivatives (Table 2).

**Table 2:** LC-MS/MS quantification of 16 target compounds from Warrigal Spinach and Kensington Pride Mango extracts.

| No | Compounds | Formula | Warrigal spinach | Kensington Pride Mango |
| --- | --- | --- | --- | --- |
| 1 | Gallic Acid | C<sub>7</sub>H<sub>6</sub>O<sub>5</sub> | 71.4 ± 1.2 | 280.6 ± 18.5 |
| 2 | *p*-hydroxybenzoic acid | C<sub>7</sub>H<sub>6</sub>O<sub>3</sub> | 122.6 ± 9.4 | 347.9 ± 47.6 |
| 3 | Cinnamic acid | C<sub>9</sub>H<sub>8</sub>O<sub>2</sub> | 449.2 ± 15.4 | ND |
| 4 | Ferulic acid | C<sub>10</sub>H<sub>10</sub>O<sub>4</sub> | 400.6 ± 44.8 | 103.21 ± 5.9 |
| 5 | Caffeic acid | C<sub>9</sub>H<sub>8</sub>O<sub>4</sub> | 259.4 ± 6.1 | ND |
| 6 | Protocatechuic acid | C<sub>7</sub>H<sub>6</sub>O<sub>4</sub> | 437.3 ± 17.3 | 103.52 ± 9.1 |
| 7 | Ellagic acid | C<sub>14</sub>H<sub>6</sub>O<sub>8</sub> | 141.1 ± 5.9 | 194.47 ± 14.5 |
| 8 | Myricetin 3-rutinoside | C<sub>27</sub>H<sub>30</sub>O<sub>17</sub> | 836.2 ± 26.9 | ND |
| 9 | *p*-coumaric acid | C<sub>9</sub>H<sub>8</sub>O<sub>3</sub> | ND | 305.3 ± 4.1 |
| 10 | Quercetin | C<sub>15</sub>H<sub>10</sub>O<sub>7</sub> | ND | 47.7 ± 1.2 |
| 11 | Mangiferin | C<sub>19</sub>H<sub>18</sub>O<sub>11</sub> | ND | 262.1 ± 20.3 |
| 12 | Pyrogallol | C<sub>6</sub>H<sub>6</sub>O<sub>3</sub> | ND | 327.60 ± 39.77 |
| 13 | Mangostinone | C<sub>23</sub>H<sub>24</sub>O<sub>5</sub> | 32.6 ± 2.4 | 104.2 ± 7.3 |
| 14 | Quinic Acid | C<sub>7</sub>H<sub>12</sub>O<sub>6</sub> | 241.4 ± 15.9 | 178.3 ± 9.6 |
| 15 | Protocatechuic acid 4-<br>*O*-glucoside | C<sub>13</sub>H<sub>16</sub>O<sub>9</sub> | ND | 39.2 ± 2.6 |
| 16 | 3,4-*O*-Dimethylgallic<br>acid | C<sub>9</sub>H<sub>10</sub>O<sub>5</sub> | ND | 142.3 ± 12.4 |

ND: not detected

Phenolic acids were the most abundant group in both samples, consistent with previous reports on Australian native plants, where phenolic acids dominate the phytochemical profile. In Warrigal spinach, cinnamic acid (449.2 ± 15.4 µg/g) and protocatechuic acid (437.3 ± 17.3 µg/g) were among the most abundant compounds, followed by ferulic acid (400.6 ± 44.8 µg/g) and caffeic acid (259.4 ± 6.1 µg/g). These results indicate a strong presence of hydroxycinnamic acid derivatives, which are known for their potent antioxidant properties. In contrast, Kensington Pride mango showed higher concentrations of *p*-hydroxybenzoic acid (347.9 ± 47.6 µg/g) and gallic acid (280.6 ± 18.5 µg/g) than Warrigal spinach, suggesting a phenolic composition dominated by hydroxybenzoic acid derivatives. Additionally, *p*-coumaric acid (305.3 ± 4.1 µg/g) was exclusively detected in mango, further highlighting compositional differences between the two matrices.

Among flavonoids, myricetin 3-rutinoside (836.2 ± 26.9 µg/g) was identified exclusively in Warrigal spinach and was among the most abundant compounds, indicating a strong contribution of flavonoid glycosides to its antioxidant capacity. In contrast, quercetin (47.7 ± 1.2 µg/g) was only detected in Kensington Pride mango, reflecting differences in flavonoid subclasses between the two samples. Other phenolic compounds, such as ellagic acid, were detected in both samples, with slightly higher concentrations in mango (194.47 ± 14.5 µg/g) compared to spinach (141.1 ± 5.9 µg/g). Similarly, quinic acid was present in both samples but showed higher levels in Warrigal spinach (241.4 ± 15.9 µg/g) than in mango (178.3 ± 9.6 µg/g). Compounds such as mangiferin (262.1 ± 20.3 µg/g) and pyrogallol (327.60 ± 39.77 µg/g) were exclusively detected in Kensington Pride mango, indicating the presence of unique bioactive metabolites in fruit matrices. Additionally, protocatechuic acid 4-O-glucoside (39.2 ± 2.6 µg/g) and 3,4-*O*-dimethylgallic acid (142.3 ± 12.4 µg/g) were only observed in mango.

Overall, Warrigal spinach exhibited higher concentrations of several key phenolic acids and flavonoid glycosides, particularly cinnamic acid, ferulic acid, caffeic acid, and myricetin derivatives, which are strongly associated with antioxidant activity. In contrast, Kensington Pride mango contained a broader diversity of phenolic compounds, including unique metabolites such as mangiferin and pyrogallol, but generally at lower concentrations than those of major antioxidant contributors. These findings are consistent with previous studies on Australian native plants, in which higher phenolic concentrations were directly associated with greater antioxidant activity. The presence of specific phenolic compounds unique to each sample further highlights the distinct phytochemical profiles and functional properties of leafy vegetables and fruit matrices.

### 3.4. Zebrafish larvae acute toxicity of Warrigal spinach and Kensington Pride mango extracts

Toxicology assessment of Warrigal spinach and Kensington Pride (Kensington Pride) mango was conducted using zebrafish embryos from 3 hpf onwards, with phenotypic analysis of each embryo every 24 hours until 96 hours. For all experiments, replicate clutches were included only if zebrafish embryos/larvae from the control group (0 mg/L) displayed normal morphology and had low mortality (<10%) throughout the experiment. Warrigal spinach showed a significant spike at 480 mg/L (Figure 2A, B). Whilst most of the mortality occurred within 24 hours of exposure, subsequent lethality was observed throughout the 96 hours of exposure, suggesting that toxicity effects accumulated. Kensington Pride Mango showed a relatively constant mortality rate across concentrations, with most mortality again occurring in the first 24 hours or by 48 hours (Figure 2C, D). This suggests that zebrafish are particularly sensitive to toxic effects during the early developmental period and that cumulative exposure to Kensington Pride Mango did not lead to such severe phenotypes once larvae had survived to 48 hpf.

**Figure 2.**
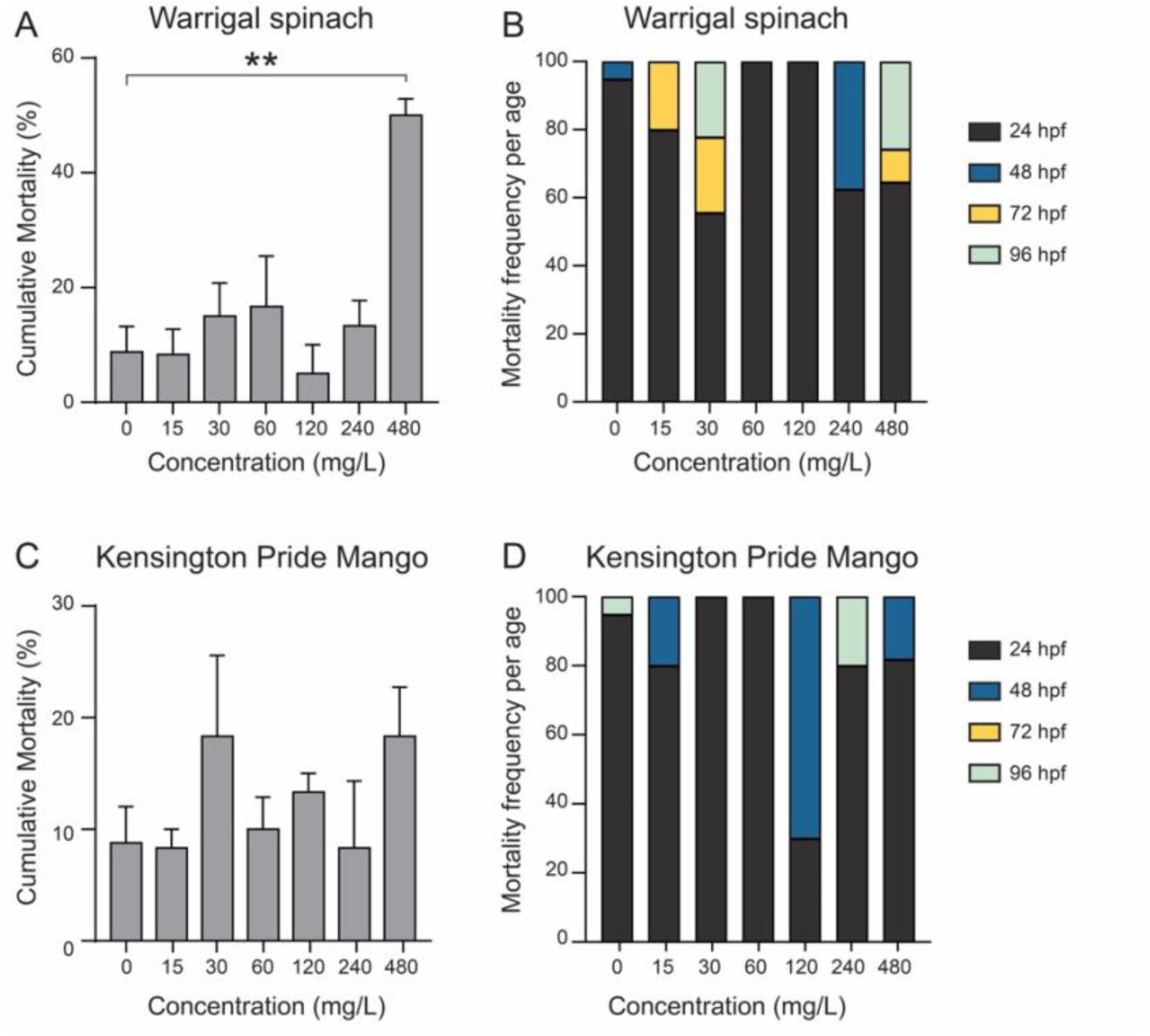
Assessment of zebrafish mortality for the whole organism exposed during 96 hours to different concentrations (0, 15, 30, 60, 120, 240 and 480 mg/L) of each plant extract. Cumulative mortality observed at 96 hours of exposure is shown for Warrigal spinach (A) and Kensington Pride mango (C). Panels B and D show the mortality frequency at exposure times of 24, 48, 72, and 96 hours. Experiments were conducted in triplicate, with 60 embryos per concentration. Data show the mean ± SEM. Asterisks represent: ** (*p* < 0.01) for Warrigal spinach (A).

The LC_50_-96h was calculated based on the cumulative percentage of the following signs of mortality: egg coagulation (0-24 h of exposure), dead embryos (24-48 h of exposure), and dead larvae (48-96 h of exposure). For Warrigal spinach extract, the LC_50_ (96 h) was 478.8 mg/L. For Kensington Pride mango extract, the LC_50_ (96 h) was >480 mg/L (our highest tested concentration). Thus, both present with a relatively high concentration range that is safe for targeted use in efficacy testing in preclinical models.

Taking a closer look at toxicological effects that did not result in lethality, we found relatively minimal effects at up to 240 mg/ml in Warrigal spinach (Figure 3A), with a sharp increase in developmental abnormalities, malformations, and mortality observed at 480 mg/ml. At this concentration, abnormalities commonly indicative of general toxicity were observed. These included significantly higher occurrence of cardiac oedema, bradycardia, yolk sac oedema, delayed yolk sac absorption and hatching delay (**** *p* = < 0.0001, and lack of blood circulation in the tail and developmental delay (*** p = < 0.001). Additionally, spine (* *p* = < 0.05) and tail (*** *p* = < 0.001) malformations were also significantly increased.

**Figure 3.**
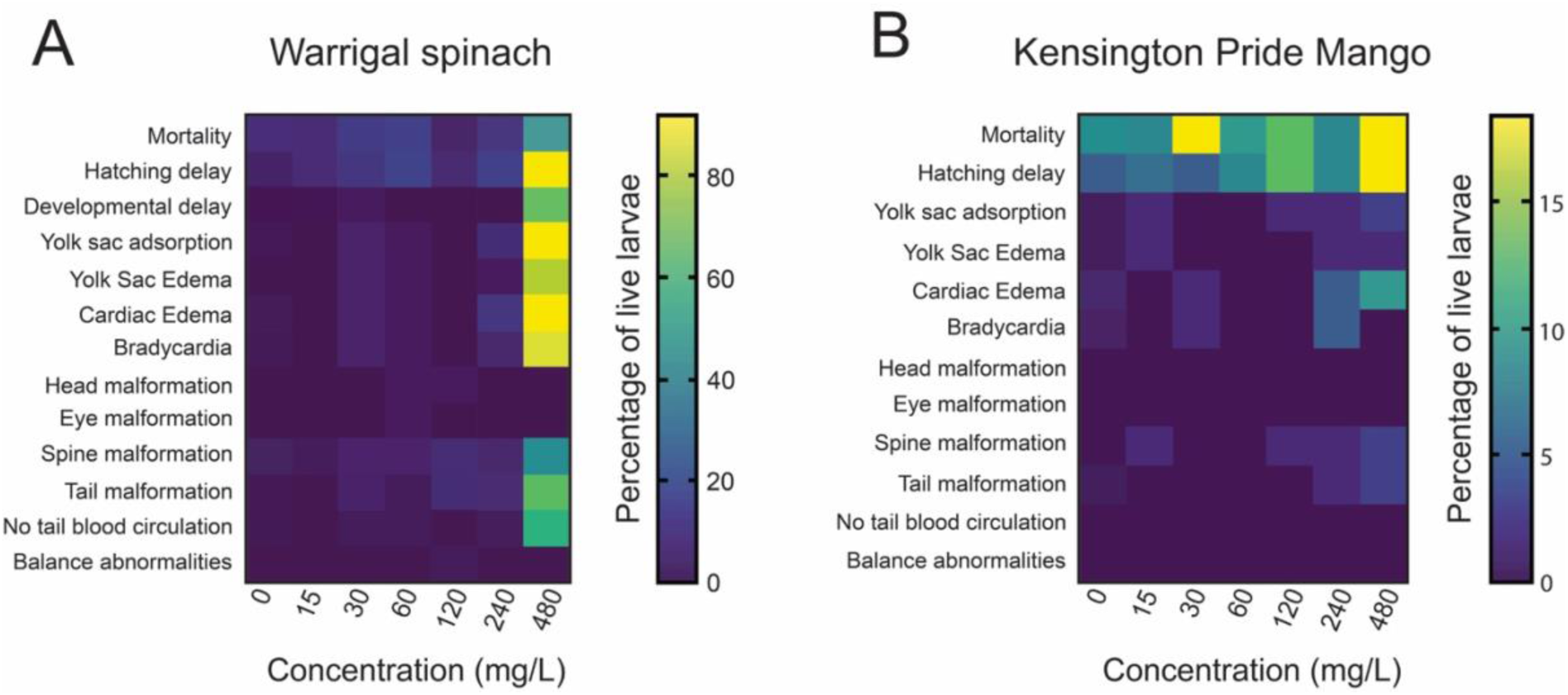
Heat maps of mortality, alterations and malformations of embryos/larvae exposed for 96 hours to the plant extracts. The mean of three replicates (n=20/replicate) is reported for Warrigal spinach (A) and Kensington Pride mango (B). Both heat maps show an increasing percentage of phenotypic traits at higher concentrations relative to controls (0 mg/L). Note that the heatmap scale differs; overall, Kensington Pride mango showed fewer toxicological signs.

For Kensington Pride mango, phenotypic indicators of toxicity were much lower overall (Figure 3B – note heatmap maximum scale is much lower). However, both mortality and hatching defects were present across all concentrations. They were also the two main phenotypes observed, even at 480 mg/ml. Other alterations, including edema, bradycardia and malformations (Figure 3B), were not statistically significantly different in frequency when compared to the control group.

Thus, while exposure to both Warrigal spinach and Kensington Pride Mango resulted in low mortality and hatching delays across various concentrations, they caused relatively few abnormalities or malformations up to at least 240 mg/L.

## 4. Discussion

As the average human lifespan is increasing, age-related health declines are becoming more prevalent and contributing to an increased burden and reduced quality of life in affected people. Oxidative stress is a highly prevalent contributor to cellular health and many of the degenerative conditions increasingly occuring in diverse organs across the body. Thus, increasing effort is being invested into understanding and leveraging the therapeutic potential of islated antioxidants and combinations of antioxidants including those found in many naturally occuring botanicals. Australia boasts an untapped resource of unique botanicals, that could be developed towards clinical use. Our work combining *in vitro* anlaysis with pre-clinical toxicology screening in the zebrafish biomedical vertebrates system allows for pre-clinical toxicology screening of natural compounds that could restore the oxidative balance and organ function.

In the present study, our characterisation showed that Warrigal spinach and Kensington Pride mango exhibited distinct phytochemical compositions and antioxidant capacities with relatively limited overlap. Warrigal spinach demonstrated significantly higher total phenolic content (TPC), total flavonoid content (TFC), and antioxidant activities (DPPH, ABTS, and FRAP) than Kensington Pride mango, indicating greater antioxidant potential. These findings are consistent with previous studies, in which higher phenolic content was strongly correlated with enhanced antioxidant activity, owing to phenolic compounds’ ability to act as hydrogen donors, reducing agents, and free radical scavengers (Muflihah et al., 2021). Similar trends have been reported in native Australian fruits such as Kakadu plum, Davidson plum, and strawberry gum and other native plants, such as bush mint and lemongrass, which showed significantly higher TPC and antioxidant activities than other fruits, with a clear positive correlation between phenolic content and DPPH and ABTS scavenging capacities (A. Ali et al., 2023; Akhtar Ali et al., 2023; Ali, Kiloni, et al., 2022; Cáceres-Vélez et al., 2022). Importantly, our findings reinforce the concept that antioxidant activity is governed not only by the presence of specific compounds but also by their overall abundance and synergistic interactions within complex phytochemical mixtures (Ilie et al., 2024).

Furthermore, the differences observed between Warrigal spinach and Kensington Pride mango may be attributed to variations in plant physiology and metabolic pathways. Leafy vegetables such as Warrigal spinach are often exposed to environmental stressors, including UV radiation, which stimulates the biosynthesis of phenolic compounds as protective metabolites. In contrast, fruit matrices such as mango may contain lower levels of phenolic antioxidants or a different composition dominated by other bioactive compounds such as carotenoids and vitamins, which may not be fully captured by the assays used in this study. Overall, these findings clearly demonstrate that Warrigal spinach is a substantially richer source of phenolic antioxidants compared to Kensington Pride mango. The strong correlation between phenolic content and antioxidant activity, supported by both the present results and previous literature, highlights the importance of phenolic compounds in determining the functional and nutritional value of plant-based foods (Ouamnina et al., 2024).

At the compound level, Warrigal spinach was characterised by higher concentrations of hydroxycinnamic acids (cinnamic, ferulic, caffeic acids) and flavonoid glycosides such as myricetin derivatives, which are well known for their potent antioxidant activity. The phenolic compound cinnamic acid has been described as a potential target for oxidative stress and inflammation (Issac et al., 2021; Pontiki & Hadjipavlou-Litina, 2018). Caffeic acid, an antioxidant in the hydroxycinnamic acid group has been identfied in tamarind (Cáceres-Vélez et al., 2022; El-Haddad et al., 2019) and has been shown protect kidneys from toxicity-induced damage in rats (Ogeturk et al., 2005), suggesting high potential for combatting oxidative stress.

The bioactive constituents and antioxidant classes in mango have been shown to vary widely depending on the geographical region, ripening stage, and which part of the mango is analysed (e.g. leaves, peel, pulp, seed coat and kernel) (Abbasi et al., 2015; Rambabu et al., 2019; Rocha Ribeiro et al., 2007). Aqueous stem bark from the related *Mangifera indica* mango has previously been shown to represent a rich source of antioxidants, contain carotenoids and polyphenolic compounds (Pam et al., 2021; Rocha Ribeiro et al., 2007; Rosalie et al., 2018). In our analysis Kensington Pride mango exhibited a distinct phytochemical profile dominated by gallic acid, *p*-hydroxybenzoic acid, mangiferin, and pyrogallol. Although compounds such as mangiferin possess strong antioxidant and anti-inflammatory properties, the comparatively lower total phenolic concentration in mango likely contributes to its reduced antioxidant capacity. Interestingly, anti-tumour / anti-cancer effects of *Mangifera indica* have also been demonstrated in mice, rats and *in vitro* colon cancer cells (Mirza et al., 2021; Rocha Ribeiro et al., 2007; Shaban et al., 2023; Shaban et al., 2022; reviewed in Yap et al., 2021). Additionally, antioxidant potential of mango pulp has been shown to be an important dietary component that may have cardioprotective effects and aid in prevention of type 2 diabetes mellitus (Fernandes & Salgado, 2016; Gondi & Prasada Rao, 2015; Tan et al., 2011b).

These findings reinforce the concept that antioxidant activity is governed not only by the presence of specific compounds but also by their overall abundance and synergistic interactions within complex phytochemical mixtures (Ilie et al., 2024).

Importantly, while Warrigal spinach demonstrated superior antioxidant capacity, both plant extracts exhibited high safety margins in the zebrafish embryonic model, indicating low acute toxicity. The LC_50_ values observed in this study (∼478.8 mg/L for Warrigal spinach and >480 mg/L for mango) classify both extracts as safe according to OECD toxicity guidelines, which categorise compounds with LC_50_ values above 100 mg/L as relatively non-toxic. These findings are consistent with previous studies (Ali, Kiloni, et al., 2022; Cáceres-Vélez et al., 2022) on native Australian fruits, where Kakadu plum and quandong peach also exhibited LC50 values greater than 480 mg/L, while Davidson plum showed moderate toxicity (LC50 ∼376 mg/L), and muntries exhibited comparatively higher toxicity (LC_50_ ∼169 mg/L).

Comparative analysis with other plant-based toxicity studies further highlights the relatively low toxicity of the extracts investigated in this study. For instance, significantly lower LC_50_ values have been reported for several medicinal plants, including turmeric (LC_50_ ∼56.67 µg/mL), Chinese motherwort (∼10–60 µg/mL), and *Euphorbia kansui* (<10 µg/mL), indicating substantially higher toxicity compared to Warrigal spinach and mango extracts (Alafiatayo et al., 2019; He et al., 2018; Zhang et al., 2018). Similarly, safflower extracts (LC_50_ ∼345.6 mg/L) and pomegranate extracts (∼196 mg/L) demonstrated moderate toxicity relative to the present findings (Wibowo et al., 2018; Xia et al., 2017). These comparisons emphasise that the extracts analysed in this study fall within a favourable safety range for potential nutraceutical or pharmaceutical applications.

Notably, previous studies have demonstrated that toxicity is not directly correlated with antioxidant capacity or total phenolic content. For example, extracts with high phenolic content have been shown to exhibit varying toxicity profiles depending on their specific phytochemical composition. In muntries, higher toxicity was attributed to compounds such as saponins, alkaloids, and estragole, which are known to exert cytotoxic, genotoxic, or hepatocarcinogenic effects. Similarly, studies on other Australian plants, such as mountain pepper, identified specific compounds (e.g., estragole and lignans) associated with increased toxicity, despite strong antioxidant activity (Ali, Kiloni, et al., 2022; Caceres-Velez et al., 2024).

These observations highlight the importance of considering both individual compounds and their interactions within complex mixtures. The toxicological effects of plant extracts are often governed by synergistic or antagonistic interactions among phytochemicals, which can enhance or mitigate the biological activity of individual constituents (Vaou et al., 2022). This complexity explains why extracts with similar antioxidant capacities may exhibit different toxicity profiles and underscores the necessity of integrating chemical characterisation with biological evaluation (Ilie et al., 2024). Furthermore, variations in toxicity across studies can also be attributed to differences in extraction methods, solvent systems, plant material, and experimental conditions. Factors such as solvent polarity, extraction time, temperature, and plant tissue type significantly influence the composition and concentration of phytochemicals, ultimately affecting both antioxidant activity and toxicity (Ali, Kiloni, et al., 2022; Altemimi et al., 2017; Kumar et al., 2017; Robbins, 2003). Therefore, direct comparisons between studies should be made cautiously, and toxicity assessments must be correlated with specific extraction conditions.

The *danio rerio* fish embryo toxicity test (FET) employed followed the standardised OECD (2013) guidelines, allowing for robust and high-throughput and cross-study comparative data for evaluating the plant-derived compounds and comprehensive evaluation of their benefit–risk profile. Owing to its genetic and physiological similarities to humans, rapid development, and transparency, zebrafish offer significant advantages over traditional mammalian models for early-stage toxicological screening of many botanicals (reviewed in Chahardehi et al., 2020; Falcao et al., 2018; Lima et al., 2025). Zebrafish are being used to assess toxicology (but not necessarily antioxidant capacity) of a growing number of ethnomedicinal plants including Java tea, bay leaf, king of bitters, chinese motherwort, ginseng, and our studies on Australian botanicals including Finger Lime, Mountain Pepper, Tamarind, Davidson plum, Kadadu, plum, Quandong peach and muntries (Alafiatayo et al., 2019; Cáceres-Vélez et al., 2022; He et al., 2018; Ismail et al., 2017; Ong et al., 2015; Wibowo et al., 2018; Xia et al., 2017; Zhang et al., 2023). Collectively, this study highlights that while phenolic-rich plant extracts such as Warrigal spinach offer enhanced antioxidant potential, their safe application depends on the complex interplay of phytochemical composition, concentration, and biological interactions, emphasising the critical need for integrated phytochemical and toxicological assessments in natural product research.

## 5. Conclusion

This study demonstrates that Warrigal spinach has significantly higher levels of phenolics and flavonoids, resulting in superior antioxidant capacity compared with Kensington Pride mango with both exhibiting diverse phytochemical profiles, dominated by flavonoids and phenolic acids. Our findings indicate that antioxidant capacity is closely associated with phenolic composition but does not directly predict toxicity, demonstrating the importance of empirical safety assessments. As additional botanicals with high antioxidant capacity and low toxicity are identified, comparative profiling of their antioxidant compositions will be essential for pinpointing specific compounds—or synergistic combinations—that confer beneficial activity without inducing adverse effects and can be prioritised for evaluation in mammalin models, which are more resource-intensive. This study highlights the importance of integrating phytochemical characterisation with *in vivo* safety assessment. Warrigal spinach emerges as a promising source of natural antioxidants, and both extracts show potential for safe application in functional foods and nutraceuticals.

## Acknowledgements

This study was supported by a J.N. Peters fellowship and K.M. Brutton Bequest from the Faculty of Science, University of Melbourne, Australia, to P.R.C-V. We are grateful to Julie Weatherhead and Anthony Hooper (owners of Peppermint Ridge Farm) for providing plant materials.

## Author contributions: CRediT

This study was conceptualised by P.R.C-V. and P.R.J.; investigation, methodology and data analysis were carried out by S.K., A.A., and P.R.C-V. The project was administered by F.D. and P.R.J., who provided the required resources within their labs. Supervision was carried out by F.D., P.R.C-V.,

J.C. and P.R.J.; original manuscript draft was written by S.K., A.A., and P.R.J. and the manuscript was revised by all authors. All authors have read and agreed to the published version of the manuscript.

## Data availability

The additional data are available in the Supplementary Materials.

## Declaration of generative AI use

During the preparation of this work, the authors used ChatGPT and Grammarly to improve grammar, structure, and flow. After using these tools, the authors reviewed and edited the content as needed and take full responsibility for the content of the published article.

## Supplementary Materials

**Figure S1.**
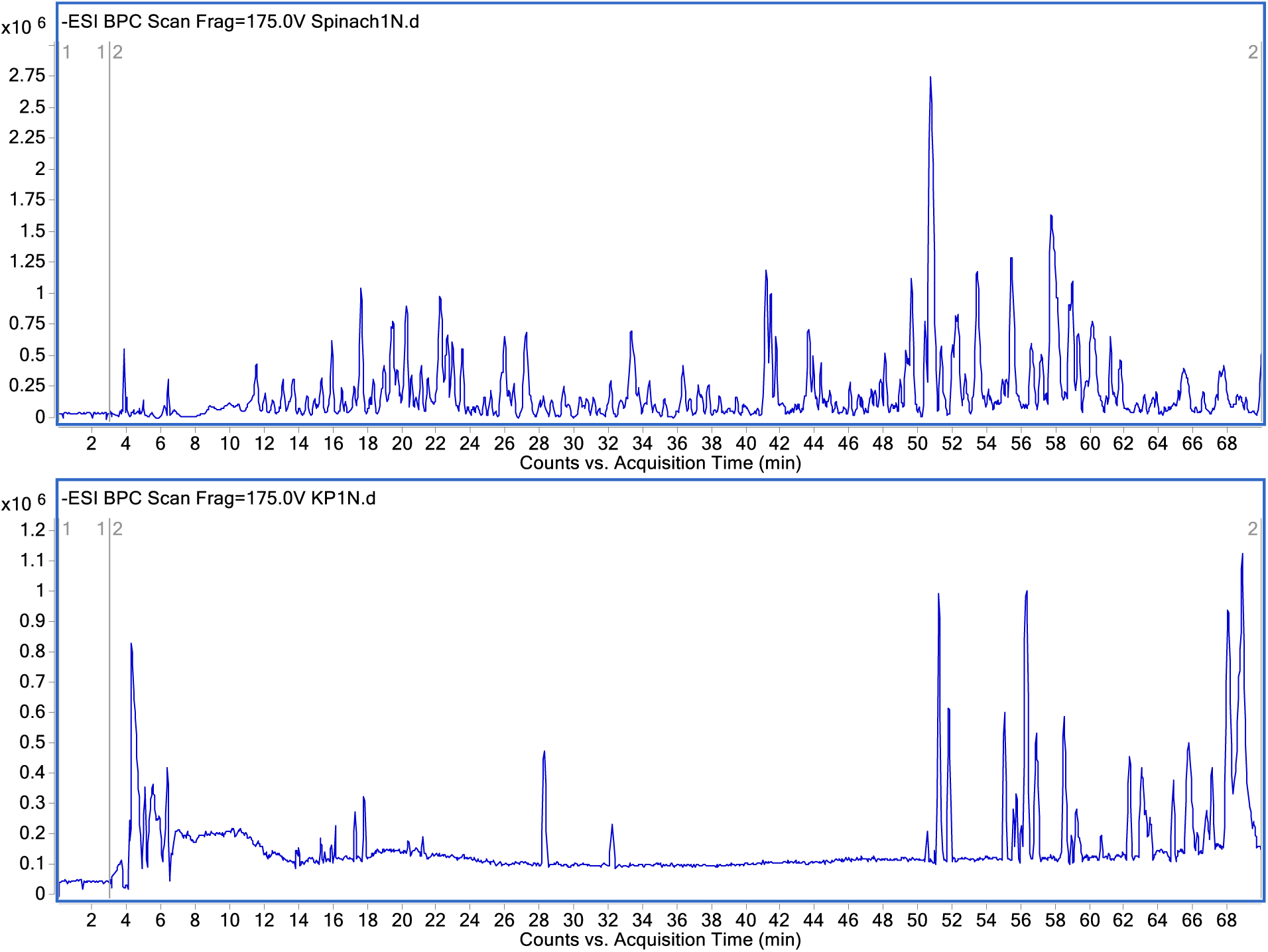
Base Peak Chromatograms (BPC) of Spinach and KP in negative mode.

**Figure S2.**
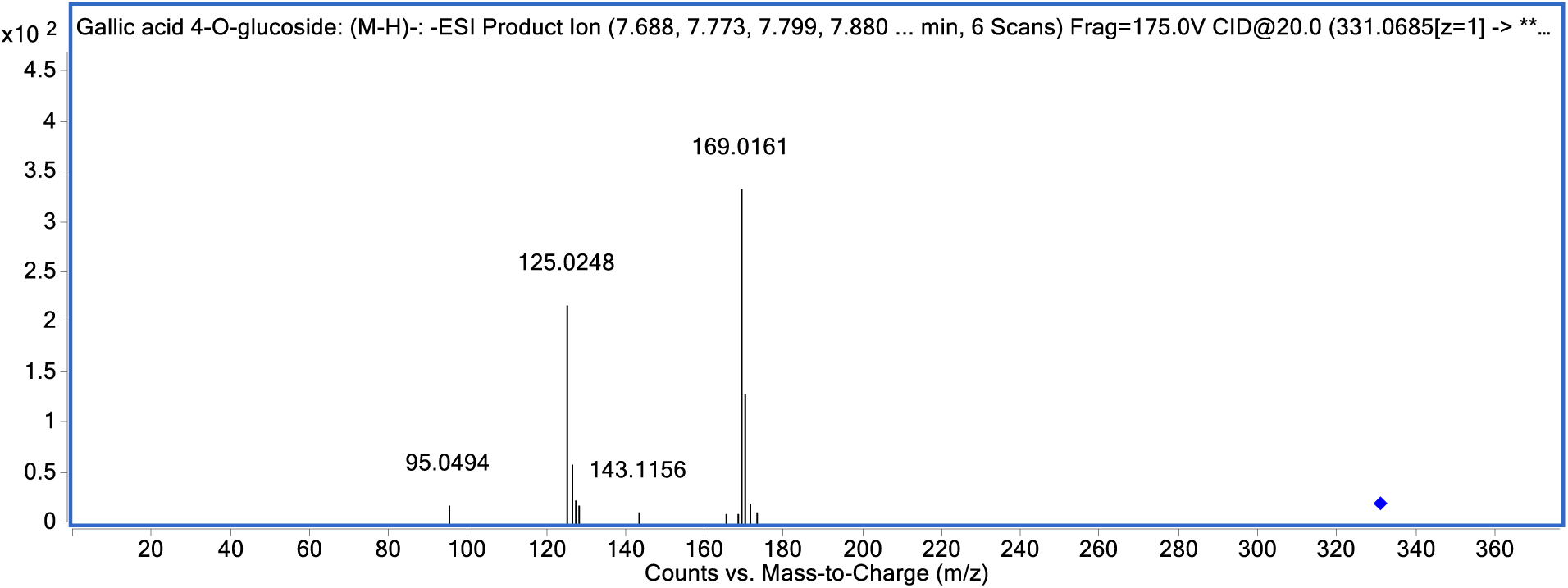
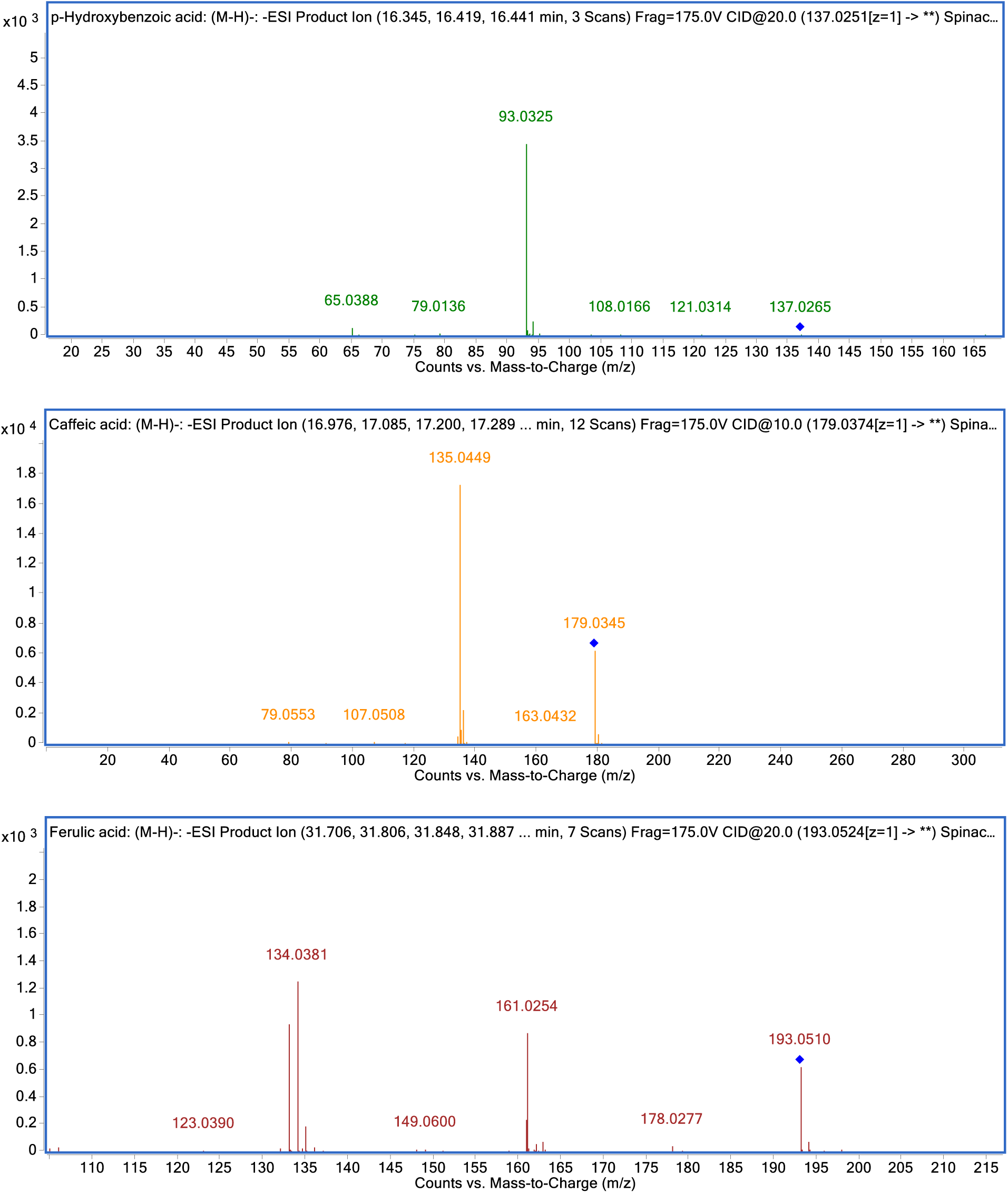
MS/MS spectra of some selected compounds: gallic acid 4-*O*-glucoside, *p*-hydroxybenzoic acid, ferulic acid, and caffeic acid.

**Table S1.**
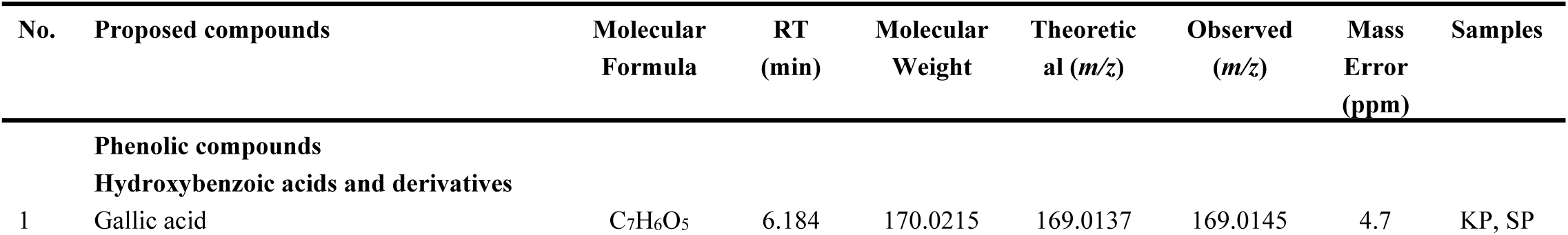
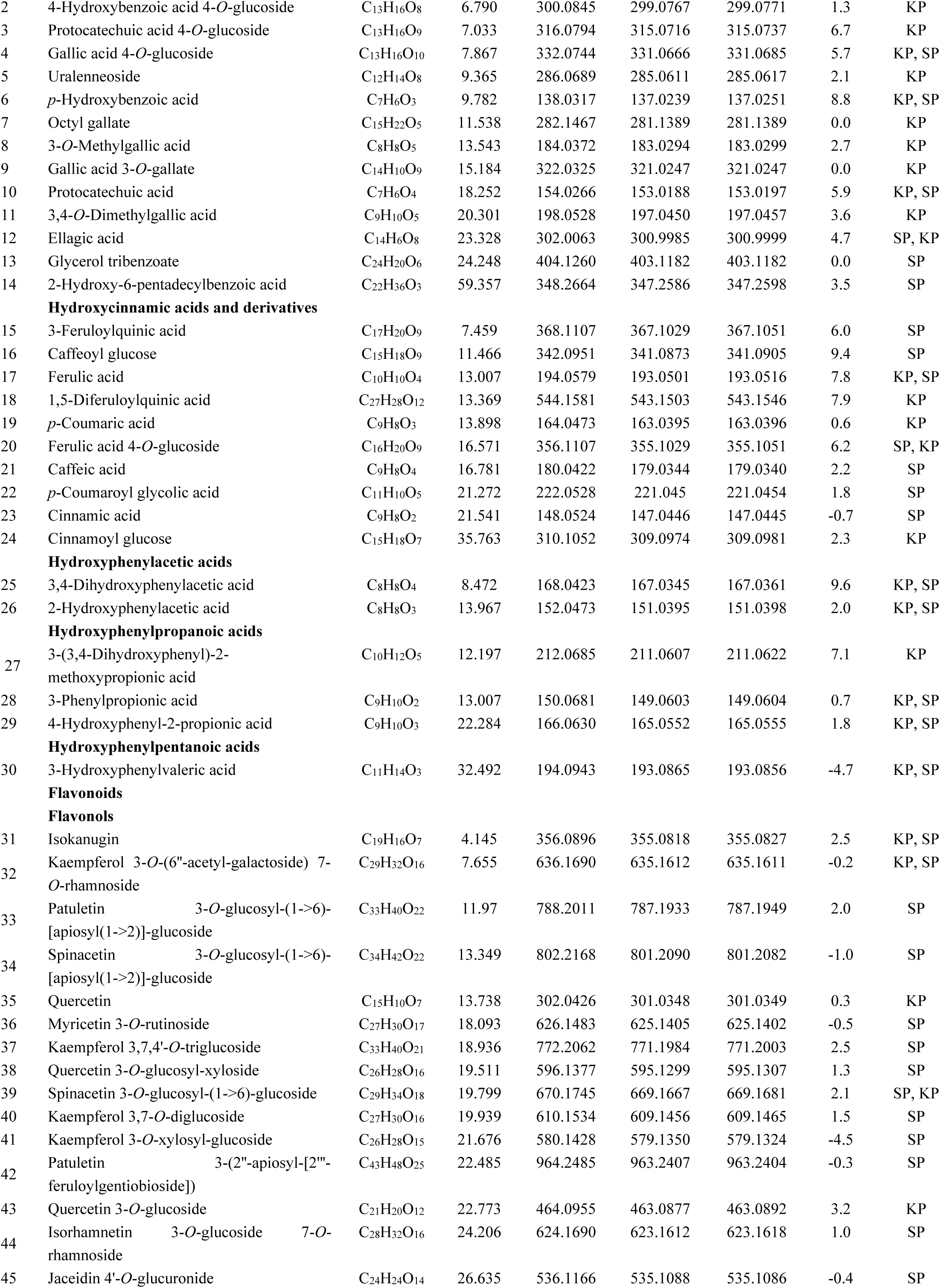
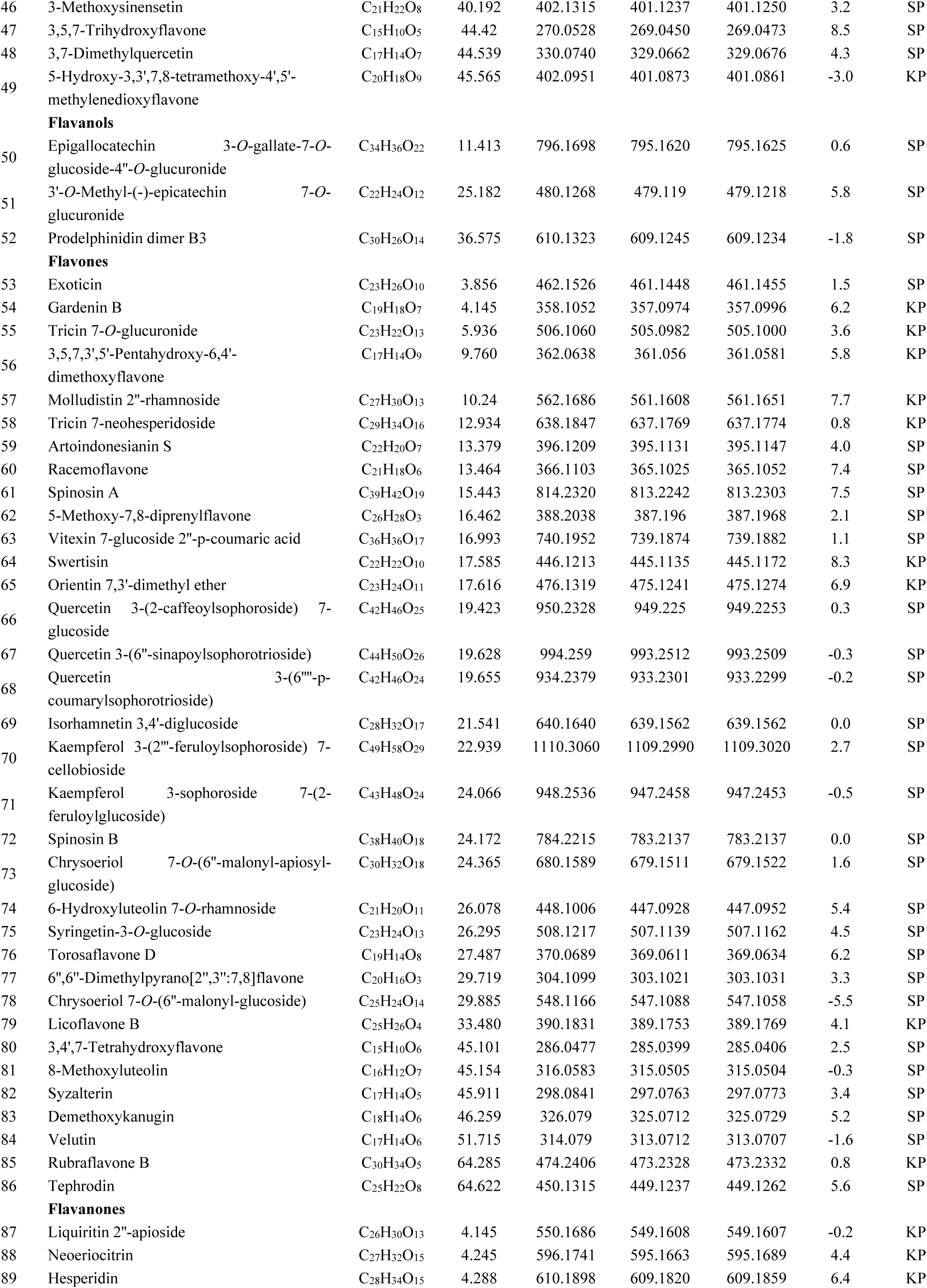
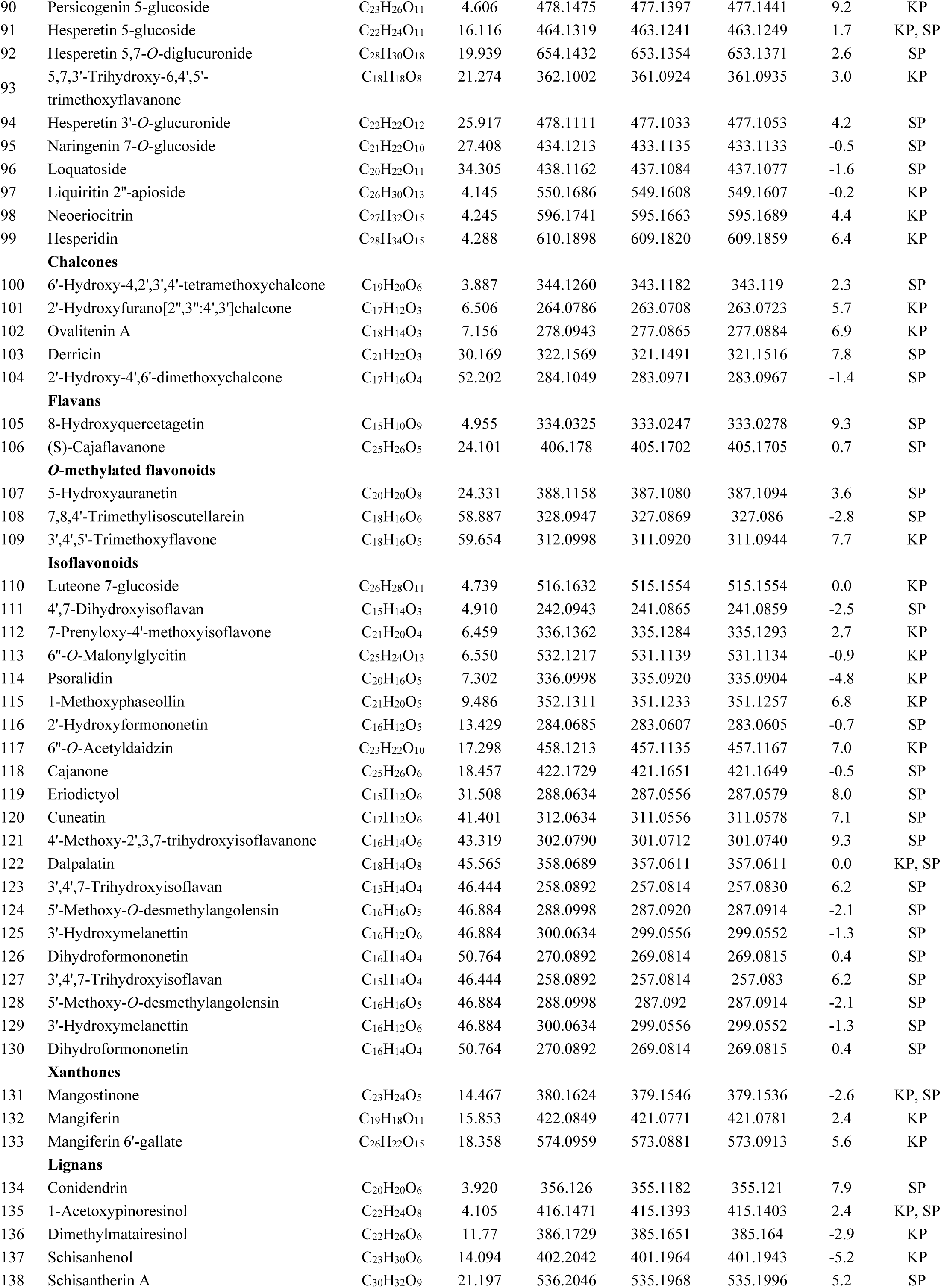
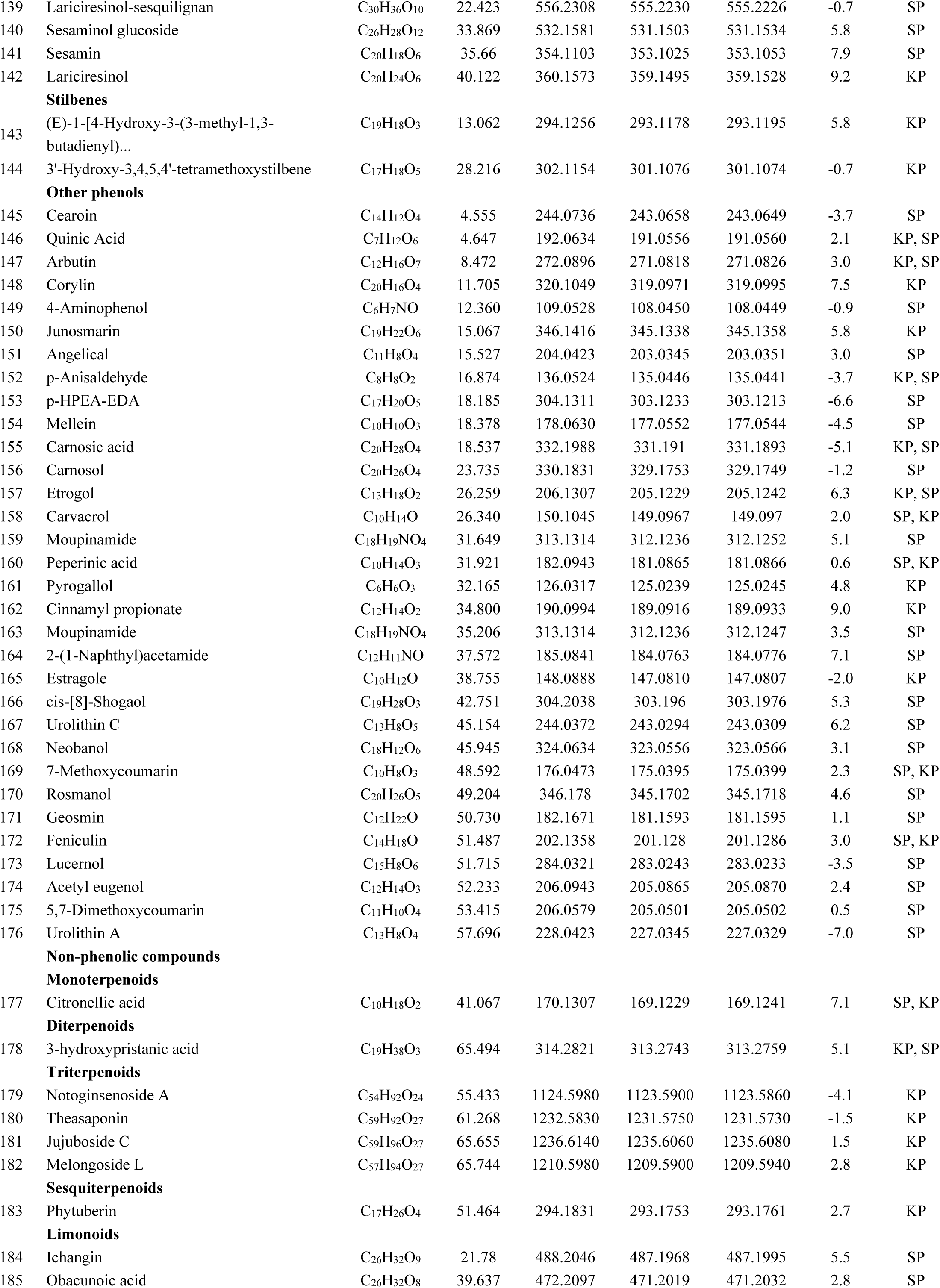
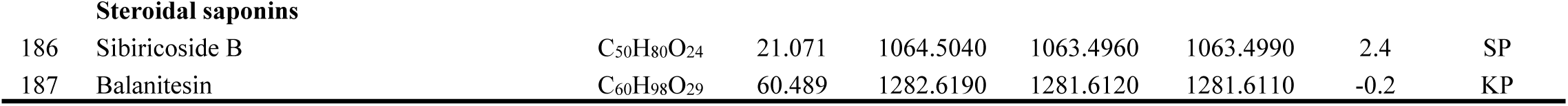
LC-ESI-QTOF-MS/MS screening and identification of phytochemicals from Spinach (SP) and Mango (KP)

